# Selective effects of a short transient environmental fluctuation on a natural population

**DOI:** 10.1101/2022.02.10.479864

**Authors:** Markus Pfenninger, Quentin Foucault, Ann-Marie Waldvogel, Barbara Feldmeyer

## Abstract

Natural populations experience continuous and often transient changes of environmental conditions. These in turn may result in fluctuating selection pressures leading to variable demographic and evolutionary population responses. Rapid adaptation as short-term response to a sudden environmental change has in several cases been attributed to polygenic traits, but the underlying genomic dynamics and architecture are poorly understood. In this study, took advantage of a natural experiment in an insect population by monitoring genome-wide allele frequencies before and after a cold snap event. Whole genome pooled sequencing of time series samples revealed ten selected haplotypes carrying ancient polymorphisms, partially with signatures of balancing selection. By constantly cold exposing genetically variable individuals in the laboratory, we could demonstrate with whole genome resequencing i) among the survivors, the same alleles rose in frequency as in the wild and ii) that the identified variants additively predicted fitness (survival time) of its bearers. Finally, by simultaneously sequencing the genome and the transcriptome of cold exposed individuals we could tentatively link some of the selected SNPs to the *cis*- and *trans*-regulation of genes and pathways known to be involved in cold response of insects, like *Cytochrome P450* and fatty acid metabolism. Altogether, our results shed light on the strength and speed of selection in natural populations and the genomic architecture of its underlying polygenic trait. Population genomic time series data thus appear as promising tool for measuring the selective tracking of fluctuating selection in natural populations.

## Introduction

Adaptation in natural populations occurs when selection acts on variable phenotypic traits with a heritable basis. There is a general agreement that selection in the wild is intense (Hoekstra et al., 2001; Kingsolver et al., 2001). It is also variable in space and time (Bell, 2010; Price, Grant, Gibbs, & Boag, 1984; Siepielski, DiBattista, Evans, & Carlson, 2011), even though there is some debate whether changes in the direction of selection are frequent or not (Kingsolver, Diamond, Siepielski, & Carlson, 2012; Kingsolver & Pfennig, 2007; Siepielski, DiBattista, & Carlson, 2009). Recent theoretical (Messer & Petrov, 2013) and empirical work (Bitter, Kapsenberg, Gattuso, & Pfister, 2019) has shown that selection in natural populations can lead to rapid adaptation, in particular of polygenic traits (Barghi, Hermisson, & Schlötterer, 2020; Jain & Stephan, 2017). If the rate of environmental change is not too fast and the population characteristics allows for effective selection, adaptation from standing genetic variation to track moving phenotypic optima is theoretically possible nearly in real-time (Matuszewski, Hermisson, & Kopp, 2015). It is therefore possible that at least some organisms, for example multivoltine species with large population sizes adaptively track their fluctuating environment (Bell, 2010).

This theoretical basis, however, has currently little empirical support from natural populations. While there are on the one hand examples of selective tracking of the fluctuating environment for phenotypic traits (de Villemereuil et al., 2020; Grant & Grant, 1989; Marrot, Garant, & Charmantier, 2017) and on the other demonstrations of rapid selectively driven changes on the molecular level (Margres et al., 2017; Yang et al., 2016; Zong, Li, & Liu, 2021), we are not aware of studies bringing together the observed temporal fitness differences among different phenotypes with the underlying molecular variants in natural populations.

In this study, we took advantage of a natural experiment to tackle the above question. We studied the genomic response of natural population of a non-biting midge to a short-term weather event, in this case a cold snap. This promised the opportunity to study the selective effects of a defined transient event as is typical for the selective regime of fluctuating environments (Bell, 2010). Non-biting midges of the Chironomid family are widely distributed aquatic insects and have a crucial role in freshwater benthic ecosystems serving as a basis of benthic food webs (Horváth, Móra, Bernáth, & Kriska, 2011; Oppold et al., 2016; Pfenninger & Nowak, 2008). *Chironomus riparius* (Meigen, 1803) is a multivoltine species with up to 15 generations per year in Europe (Oppold et al., 2016). Therefore, the different generations are subjected to widely varying environmental conditions. Accordingly, extensive research on temperature and photoperiod has shown that several traits can and do adapt locally (Waldvogel et al., 2018), and temporally among seasons (Doria, Caliendo, Gerber, & Pfenninger, 2022; Foucault, Wieser, Waldvogel, Feldmeyer, & Pfenninger, 2018). But also other factors are known to act as selection pressures on this species (e.g. organic load, Kraak *et al*. (2000), conductivity, (Pfenninger & Nowak 2008), nitrogen, (Nemec *et al*. 2012), temperature, (Nemec *et al*. 2013) and anthropogenic substances, (Nowak *et al*. 2009). The high effective and demographic population size (> 1,000,000, Waldvogel *et al*. (2018)) and the very high number of offspring per breeding pair (400-800) allows for rapid adaptation (Pfenninger & Foucault, 2020a). Since genomic resources and parameters are available (Schmidt *et al*. 2020; Oppold & Pfenninger 2017) and the species is amenable for evolutionary experiments in the laboratory (Foucault, Wieser, Waldvogel, & Pfenninger, 2019), the species is increasingly becoming a model for molecular ecology and the emerging field of evolutionary ecotoxicology (Doria, Hannappel, & Pfenninger, 2022; Doria, Waldvogel, & Pfenninger, 2021).

In this study we focussed on the following research questions:

- Does normal, transient environmental variation like a cold snap trigger measurable molecular selection in a natural population?
- Are the putatively selected SNP-loci linked to longer survival also under experimental cold exposure conditions?
- Can we link the identified variants to lower level phenotypic changes, i.e. gene expression differences?

## Material and Methods

### Temporal sampling of natural population

In the course of routine sampling for another project (Pfenninger & Foucault, 2020b), we sampled larvae of the species *Chironomus riparius* on Feb. 15 2018 with a sieve at a single site situated in a small river (Hasselbach, Hessen, Germany 50.167562°N, 9.083542°E) following the protocol of Foucault *et al*. (2019b). The sampling site is located close to a wastewater treatment plant (Abwasserverband Freigericht) that continuously monitors physical and chemical water parameters, which they generously provided. A few days after the sampling, the air temperature in the region fell substantially below zero for a couple of days, which eventually drove the water temperatures at the sampling site from the long-term average of 9-10°C during this time of the year down to about 5°C for 2 consecutive days (Figure 1a). We seized the opportunity to obtain another sample of 80 individuals from the same site. Please note that no reproduction takes place in this species at temperatures below ∼12-14°C and thus the same generation was sampled. A third sample from the same site was obtained in September 2018, about 6-7 generations later (Oppold et al., 2016). The taxonomic identity of the larvae was ascertained by DNA-barcoding of a mitochondrial (COI) and a nuclear locus (L44). Eighty thus identified *C. riparius* were pooled and subjected to Pool-sequencing (see below).

**Figure 1.**
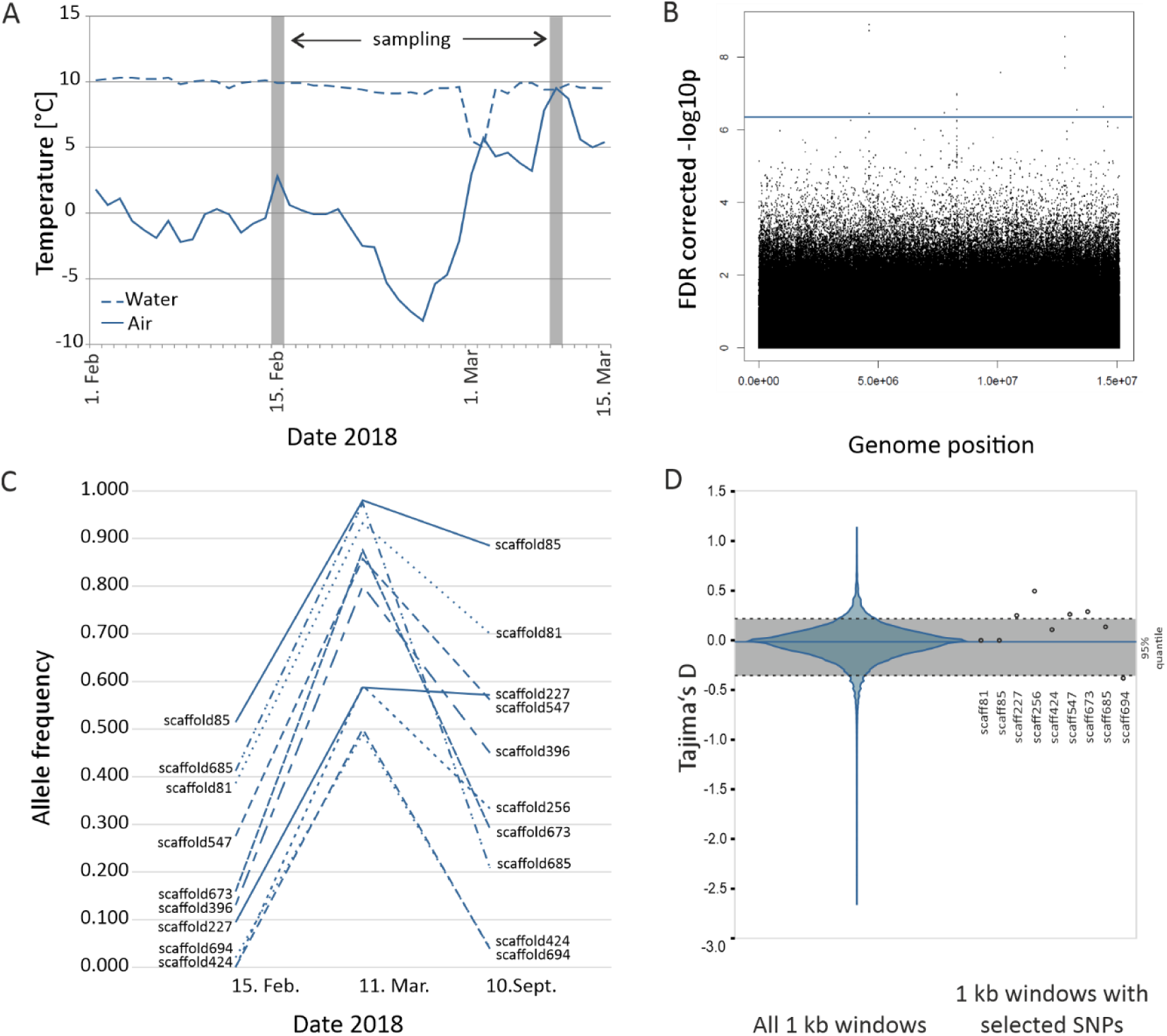
Field data. A) Air and water temperature curves at the sampling site with sampling dates (grey bars). B) Manhattan plot of the genome-wide SNPs FDR-corrected -log_10_p values, contrasting the population pools sampled before and after the cold snap. The horizontal line shows the inferred threshold. C) Allele frequency trajectories at potentially selected loci before and after the cold snap and 6-7 generations later in September 2018. D) Left: Violin-plot of the distribution of Tajima’s D for all 177,185 1 kb windows in the genome The mean is indicated by a blue horizontal line, the grey area indicates the 90% quantile around the mean. The dashed lines mark the beginning of the upper, respectively lower 5% quantile. Right: Tajima’s D for the 1kb windows harbouring the putatively selected SNPs. For one window (scaff396) Tajima’s D could not be computed.

### Population genomic analyses

DNA was extracted for the three pools from the field using the Quiagen blood and tissue extraction kit on pooled samples of 80 larval head capsules, respectively. Integrity and quality of extracted DNA was controlled using electrophoresis, and the DNA concentration for each samples measured with a Qubit fluorimeter (Invitrogen).

Whole genome pool-sequencing was carried out on an Illumina MiSeq with 250bp paired end reads. Reads were trimmed using the wrapper tool Autotrim (Waldvogel *et al*. 2018) that integrates Trimmomatic (Bolger *et al*. 2014) for trimming and FastQC (Andrews 2010) for quality control. The trimmed reads were then mapped on the latest *C. riparius* reference genome (Schmidt *et al*. 2020) using the BWA mem algorithm (Li & Durbin 2009). Low quality reads were subsequently filtered and SNPs were initially called using Samtools (Li *et al*. 2009). The pipelines PoPoolation1 v.1.2.2 and PoPoolation2 v.1.201 (Kofler *et al*. 2011a; Kofler *et al*. 2011b) were used to call SNPs and remove indels. Allele frequencies for all SNPs with coverage between 15x and 70x were estimated with the R library PoolSeq (Taus *et al*. 2017).

Selected SNP loci were identified by their allele-frequency change (AFC) larger than expected by sampling variance. Neutral simulations were used to compute false discovery rate q-values < 0.001 with parameters (number of SNPs, starting allele frequencies matching the ancestral population, sequence coverage, number of generations) matching those of the respective samples. To be conservative, we calculated the drift for one generational passage. As effective population size, we used 15,000, which constitutes a very conservative estimate as well (see Waldvogel *et al*. (2018)). All calculations and simulations were performed with the R-library poolSeq (Taus *et al*. 2017).

We used Popoolation1 to calculate Tajima’s D for all non-overlapping 1 kb windows in the genome. This window size was chosen based on the short average LD (< 150 bp) in this species (Pfenninger & Foucault, 2020b).

### Experimental confirmation

To verify the association of the SNPs to survival under cold stress, we exposed 160 4^th^ instar larvae from a laboratory population initially gained from the same population (Pfenninger & Foucault, 2020b) to 4°C until they died or survived for at least 28 days. The individuals were kept separately in 2 cm well plates with at least 1 cm water column in a normal fridge. We checked daily whether they were still alive by touching them to see if they still moved. Dead larvae or larvae still alive on the 28^th^ day were individually transferred to tubes filled with 70% alcohol and the day of their death recorded. Of these we chose 30 individuals which died early and 30 individuals which died late or even survived until the end of the 28^th^ days for resequencing.

To link genotype to gene expression we were not able to use the dead individuals from the survival experiment mentioned above. We therefore performed a corresponding short-term experiment exposing another set of 54 4^th^ instar larvae from the same laboratory population to 4°C, this time for three days only in order to guarantee for survival. After these three days, 36 living individuals were cut into three pieces on a −80°C cool pad. We cut two segments from a mid-body segment from each larva for subsequent DNA isolation and resequencing, and the rest of the individual was stored at - 80°C for later RNA-isolation.

### DNA/RNA isolation and sequencing

In total, DNA was isolated from 96 individuals (60 from the long-term, and 36 from the short-term 4°C exposure experiments) using the Qiagen® blood&tissue kit. RNA was extracted using the Quick-RNA Miniprep kit (Zymo Research). Library preparation and 150bp paired-end sequencing was conducted on a NovaSeq platform.

### Identification of individual genotypes

Quality trimming and mapping of reads was conducted similar to the approach outlined above. Genotypes at the SNP positions identified in the PoolSeq approach were called with bcftools v.1.10.2 (Li, 2011). The genotypes were cross-checked manually for a random sub-sample of individuals with IGV viewer v.2.8.2 (Thorvaldsdóttir, Robinson, & Mesirov, 2013). We calculated the mean number of potentially adaptive alleles (i.e. those that rose in frequency in the natural population) per variable locus (MNAA) for each resequenced individual as quantitative measure of the multi-locus genotype at the respective loci, thereby assuming an additive genotype-to-phenotype relationship (Sella & Barton, 2019).

### RNA-Seq analysis of cold-exposed individuals and co-expression networks

Adapters were trimmed and quality checked with TrimGalore (Krueger, 2016). HiSat2 v.2.1.0 (Kim, Paggi, Park, Bennett, & Salzberg, 2019) was used to map the reads to the *C. riparius* genome (Schmidt et al. 2020). The counts table was created with HTSeq (Anders, Pyl, & Huber, 2015). To prevent spurious results due to low read counts, we removed genes with less than 10 reads in at least four samples, and samples C4 and C6 due to missing allele frequency information. The differential gene expression analysis was conducted using DESeq2 (Love, Huber, & Anders, 2014) with mean allele frequency as continuous variable*(Love et al*., *2014)(Love et al. 2014)*.

To identify networks of co-expressed genes (modules), we constructed a weighted gene co-expression network analysis using the R package WGCNA (Langfelder & Horvath, 2008), based on the genes that had passed the quality filtering step for the expression analysis (N = 8,264). Gene counts were normalized using the *varianceStabilizingTransformation* function from DESeq2 (Love et al., 2014). Following the WGCNA guidelines, we picked a soft-thresholding power of 6 for adjacency calculation. To associate modules to mean allele frequencies, we first calculated the modules’ eigengene using the *moduleEigengenes* function and tested for module trait correlation using the *corPvalueStudent* function. To obtain up-to-date annotations, we ran a local blastp (Altschul, Gish, Miller, Myers, & Lipman, 2008) of the *C. riparius* proteins versus the non-redundant protein database (version January 2022). We ran Interproscan v.5.53-87.0 (Jones et al., 2014) locally to obtain GO information using the *C. riparius* predicted proteome. The GO enrichment analysis on genes within the significant module was performed with the R package TopGO (Alexa & Rahnenführer, 2016), using the ‘parentchild’ algorithm and the Fishers exact test for significance.

### Association of transcription rates with genotypes

We used a Bayesian approach (Bååth, 2014) to test for association of genotypes at the identified variable SNP positions and normalised gene expression for genes within +/- 200 kb on the same scaffold.

## Results

### Cold snap and Sampling

From the 21^st^ of Feb. 2018, the air temperature at the sampling site dropped for 10 consecutive days below zero, with a minimum daily average of −8°C on the 27^th^ of February 2018. The water temperature, usually fluctuating around 10°C in winter, started to fall slowly as well. At the end of this period, the water temperature dropped steeply to 5°C for two consecutive days (Figure 1A).

### Large allele frequency changes within a single generation

The allele frequency changes of 19 SNPs between before and after the cold snap could not be explained by sampling variance (Figure 1B). These SNPs were therefore considered as candidates for selection (Table 1). Some of these SNPs occurred on the same scaffold in close spatial proximity to other such SNPs (within 40-750 bp, Table 1). As resequencing data showed, the rising alleles at these SNPs were linked to haplotypes (data not shown). We considered the regions with several SNPs therefore as a single locus and the SNP with the largest AFC was used as marker SNP for these linked haplotypes. Taking this into account, ten loci, each on a different scaffold of the reference genome and thus most likely physically unlinked (Pfenninger & Foucault, 2020b), were potentially affected by selection. We refer to these loci by their scaffold numbers hereafter (e.g. scaffold227).

**Table 1.** Outlier SNPs in field data.

| Scaffold | position | reference allele | alternate allele | rising allele | start allele freq. | end allele freq. | AFC | $-\log_{10}p$ | p fdr-corrected | Tajima's D<br>1k |
| --- | --- | --- | --- | --- | --- | --- | --- | --- | --- | --- |
| scaffold81 | 616771 | T | C | T/ref | 0.386 | 0.932 | 0.545 | 6.636 | 0.035 | -0.0050318 |
| scaffold85 | 970786 | G | C | G/ref | 0.514 | 0.980 | 0.467 | 6.224 | 0.046 | -0.0162351 |
| scaffold227 | 185498 | A | G | G/alt | 0.094 | 0.587 | 0.493 | 6.267 | 0.046 | 0.2564445 |
| scaffold256 | 10257 | C | G | G/alt | 0.000 | 0.591 | 0.591 | 8.734 | 0.001 | 0.4939933 |
| ~750 bp | 10279 | A | G | G/alt | 0.000 | 0.765 | 0.765 | 8.897 | 0.001 |  |
|  | 11003 | G | C | C/alt | 0.086 | 0.867 | 0.781 | 6.455 | 0.037 |  |
| scaffold396 | 51484 | A | G | A/ref | 0.130 | 0.800 | 0.670 | 6.466 | 0.037 | NA |
| scaffold424 | 450404 | C | G | G/alt | 0.000 | 0.557 | 0.557 | 6.266 | 0.046 | 0.1154956 |
| ~400 bp | 450407 | G | T | T/alt | 0.000 | 0.557 | 0.557 | 6.266 | 0.046 |  |
|  | 450623 | G | A | A/alt | 0.000 | 0.500 | 0.500 | 6.561 | 0.035 |  |
|  | 450627 | A | C | C/alt | 0.000 | 0.534 | 0.534 | 6.967 | 0.019 |  |
|  | 450663 | C | T | T/alt | 0.000 | 0.500 | 0.500 | 6.236 | 0.046 |  |
|  | 450800 | C | T | T/alt | 0.000 | 0.533 | 0.533 | 6.993 | 0.019 |  |
| scaffold547 | 301068 | T | C | T/ref | 0.275 | 0.857 | 0.582 | 7.582 | 0.006 | 0.2763451 |
| scaffold673 | 15211 | G | T | T/alt | 0.140 | 0.857 | 0.718 | 8.015 | 0.003 | 0.3208233 |
| ~40 bp | 15229 | A | G | G/alt | 0.163 | 0.862 | 0.699 | 7.706 | 0.005 |  |
|  | 15257 | C | T | T/alt | 0.159 | 0.875 | 0.716 | 8.569 | 0.001 |  |
| scaffold685 | 402992 | T | C | T/ref | 0.412 | 0.974 | 0.562 | 6.190 | 0.047 | 0.1319014 |
| scaffold694 | 289921 | T | G | T/ref | 0.020 | 0.489 | 0.469 | 6.552 | 0.035 | -0.3782241 |

The starting frequencies of the rising alleles at these loci before the cold snap were highly variable, spanning the range from undetectably rare (scaffold424 and scaffold674) to the majority allele (scaffold85, Figure 1C). All candidate alleles rose in frequency by at least about 0.5 (0.467-0.781, Table 1, Figure 1C). In September 2018, the allele frequencies of all but one locus (scaffold227) dropped back towards the level they had before the cold snap event (Figure 1C, Table 3).

The effect of the cold snap on allele frequency spectra varied between candidate loci. Comparing Tajima’s D in the 1 kb windows encompassing the selected regions with the distribution of all 1kb windows showed that four selected regions had a Tajima’s D in the upper 5% quantile (scaffold227, scaffold256, scaffold547, scaffold673), indicating balancing selection. One value (scaffold694) was in the lower 5% tail, suggesting a recent selective sweep. The remaining values were inconspicuous (four) or not calculable (one, Figure 1D).

### Validation experiment

Of the 160 larvae constantly exposed to 4°C, the first larvae died after 15 days. Mortality on day 21 was extraordinarily high (53 individuals, 34%). After 28 days, 17 (11%) larvae were still alive. For seven individuals, the dying day could not be clearly determined. The survival distribution can be found in Supplemental Figure 1.

### Genotyping of experimental individuals

For 59 of the 60 randomly selected individuals from the experiment, resequencing was successful. The individuals could be genotyped on average at 9.02 out of the 10 loci. Two loci were fixed for the rising allele in the experimental sample (Table 2) and were thus not further considered.

**Table 2.** Nearest genes to potentially selected loci.

| Scaffold | position | distance to closest gene | gene start | gene end | annotation | Annotation spatially most proximate gene |
| --- | --- | --- | --- | --- | --- | --- |
| scaffold81 | 616771 | 5044 | 621815 | 629417 | scaffold81-snap-gene-1.377 |  |
| scaffold85 | 970786 | 120 | 970906 | 973122 | scaffold85-snap-gene-1.251 | uncharacterized protein LOC119074020 |
| scaffold227 | 185498 | 6810 | 194991 | 196989 | scaffold227-augustus-gene-0.165 | cytochrome P450 28a5 |
| scaffold256 | 10257 | 1802 | 6914 | 8455 | scaffold256-snap-gene-0.15 | delta-aminolevulinic acid dehydratase-like |
| scaffold396 | 51484 | 24716 | 12088 | 26768 | scaffold396-processed-gene-0.5 | trichohyalin isoform X3 |
| scaffold424 | 450800 | 265 | 451065 | 456420 | scaffold424-augustus-gene-0.206 | uncharacterized protein LOC109541342 isoform X2 |
| scaffold547 | 301068 | 1357 | 297539 | 302425 | scaffold547-processed-gene-0.68 | protein dachsous |
| scaffold673 | 15257 | 208 | 15465 | 17951 | scaffold673-augustus-gene-0.265 | histone acetyltransferase p300 |
| scaffold685 | 402992 | 42771 | 358566 | 360221 | scaffold685-processed-gene-0.94 | - |
| scaffold694 | 289921 | 16169 | 306317 | 308300 | scaffold694-snap-gene-0.260 | - |

In the 59 individuals used in the experiment, all candidate alleles had a considerably higher start frequency than in the natural population before the cold snap (Table 3). Nevertheless, among the survivors at day 25, the frequency of all candidate alleles rose between 0.04 and 0.22 in the course of the experiment with moderate (60.6%) to very high (98.4%) posterior probability (Figure 2A). The mean number of potentially adaptive alleles per variable locus (MNAA) locus ranged between 0.50 and 1.38 among the individuals in the experiment with a mean of 0.98 (s.d. 0.18). The correlation of this value with the length of survival was positive with almost certainty (posterior probability 99.3%). The association between the variables was moderately strong (most likely estimate of r = 0.33, 95% high density interval between 0.08 and 0.55, Figure 2B).

**Table 3.** Allele frequencies in the experiment. AF = allele frequency.

| Scaffold | position | reference allele | alternate allele | rising allele | AF Feb18 | AF Mar18 | AF Sep18 | AF in exp pop |
| --- | --- | --- | --- | --- | --- | --- | --- | --- |
| scaffold81 | 616771 | T | C | T/ref | 0.386 | 0.932 | 0.700 | 1.000 |
| scaffold85 | 970786 | G | C | G/ref | 0.514 | 0.980 | 0.885 | 0.907 |
| scaffold227 | 185498 | A | G | G/alt | 0.094 | 0.587 | 0.571 | 0.339 |
| scaffold256 | 10257 | C | G | G/alt | 0.000 | 0.591 | 0.333 | 1.000 |
| scaffold396 | 51484 | A | G | A/ref | 0.130 | 0.800 | 0.450 | 0.894 |
| scaffold424 | 450623 | G | A | A/alt | 0.000 | 0.500 | 0.038 | 0.724 |
| scaffold547 | 301068 | T | C | T/ref | 0.275 | 0.857 | 0.561 | 0.839 |
| scaffold673 | 15257 | C | T | T/alt | 0.159 | 0.875 | 0.291 | 0.902 |
| scaffold685 | 402992 | T | C | T/ref | 0.412 | 0.974 | 0.208 | 0.991 |
| scaffold694 | 289921 | T | G | T/ref | 0.020 | 0.489 | 0.041 | 0.414 |

**Figure 2.**
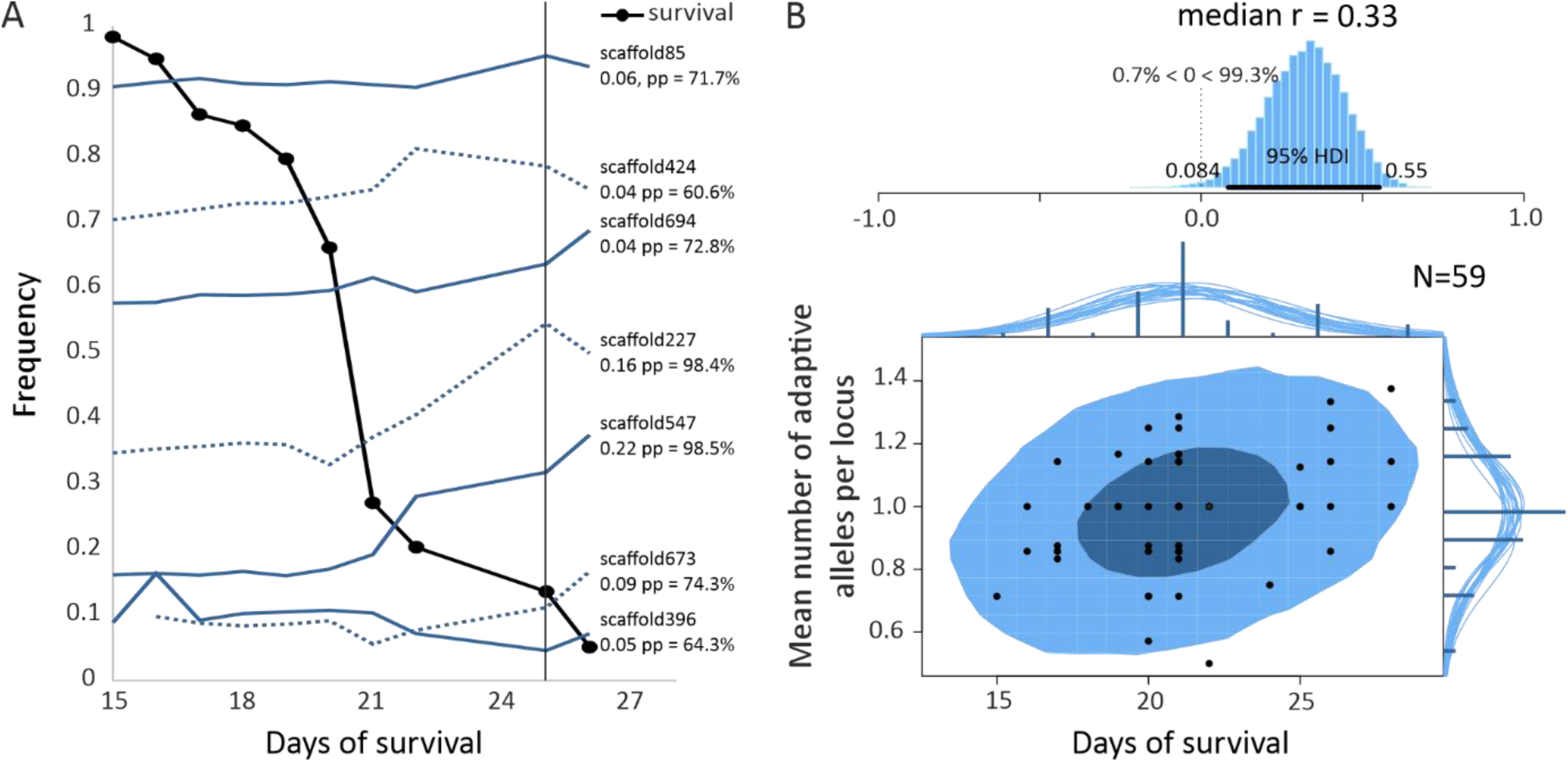
Experimental data. A) Temporal course of allele frequency changes at polymorphic candidate loci and survival of individuals during the experiment. Shown in blue are the frequency trajectories of the selected alleles in the natural population. The values to the right show the median Bayesian estimate of increase and the associated posterior probability of this value being larger than zero for the respective alleles. B) Bayesian estimate of correlation between the mean number of potentially adaptive alleles per locus and the survival time.

### Genotype associations with local and global gene expression data

For 31 individuals, we obtained gene expression data for 11,386 of the 13,449 annotated genes (85%). Only two loci showed all possible genotypes at the selected loci in the individuals used for transcription analysis (scaffold 227 and scaffold 694). On their scaffolds, the genotypes were strongly associated with the expression levels of one (scaffold 227, Figure 3A), respectively two (scaffold 694, Figure 3B) genes. On scaffold 227, the selected SNP was in an exon of the gene to which is was associated via the genotype specific expression levels. Specifically, the selected allele was associated to a higher expression rate. The gene (scaffold227_gene0.177) is annotated as Cytochrome P450, family 6 (CYP6). The two genes with expression levels strongly associated to genotypes on scaffold 694 were both roughly 100 kb away from the selected SNP. The first gene, scaffold694_gene0.209 codes for protein (RFT1) that is essential for protein N-glycosylation and also here the positively selected allele was associated with an increase of transcription. In contrast, for the second gene scaffold694_gene0.209, a transmembrane receptor protein tyrosine phosphatase (DEP1), a lower transcription level was associated with the rising allele.

**Figure 3.**
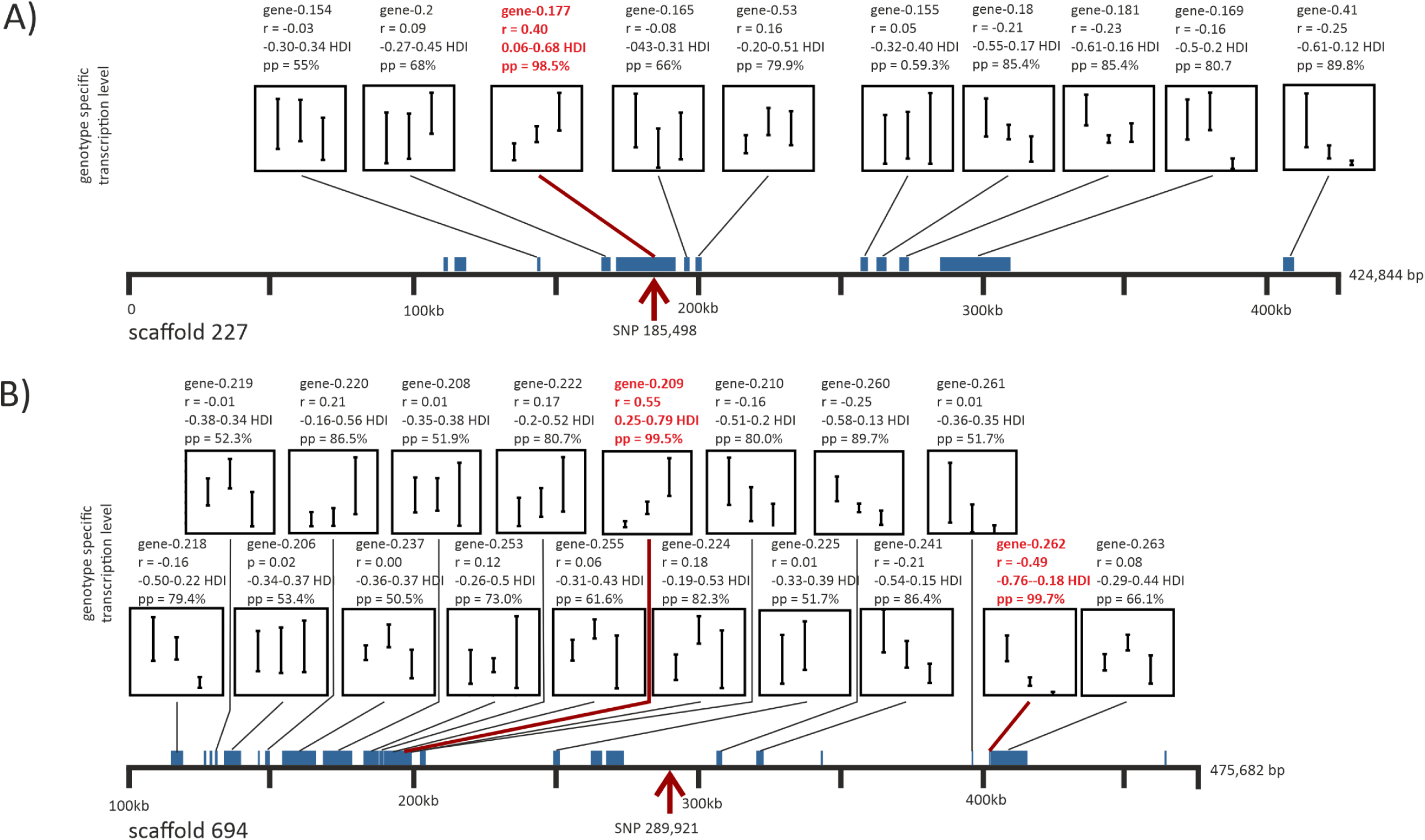
Associations of genotype at selected sites with gene expression levels of genes within +/- 200 kb on the same scaffold. The position of the selected SNP is indicated with a red arrow. Annotated genes are indicated as blue bars, whose length is proportional to the length of the gene. For genes with expression data available, the genotype specific transcription level is given in a plot. Within plot, the bars represent the standard deviation range of gene expression variation for the possible genotypes, from left to right: homozygous falling allele, heterozygous, homozygous rising allele. Above the panels, the Bayesian statistics for association are given. r = coefficient of association, HDI = 95% high density interval, pp = posterior probability for the association coefficient being larger or smaller than zero, respectively. Highlighted in red are genes with a HDI not comprising zero and pp > 95%. A) scaffold 227 with selected SNP at position 185,498 with a upregulation associated with the rising allele at gene 0.177. The SNP is positioned in an exon of the gene, B) scaffold 694 with selected SNP at position 289,921. One gene (0.209), situated about 100 kb from the SNP showed a strong upregulation associated with the rising allele. Another gene (0.262), more than 100 kb away showed a strong downregulation.

In total, we found the expression of 28 genes to be significantly associated with the MNAA. Nine out of these (32%) belong to genes of (larval) cuticule proteins, endocuticule proteins or endochitinase. One of the 20 distinct co-expressed modules identified, module-cyan (containing 169 genes), covaried substantially with MNAA (Figure 4A, r = 0.41, p=0.03). A GO-enrichment analysis revealed that gene functions related to fatty acid metabolism were overrepresented in this module (Figure 4b). The cold snap candidate loci themselves were not part of module cyan.

**Figure 4.**
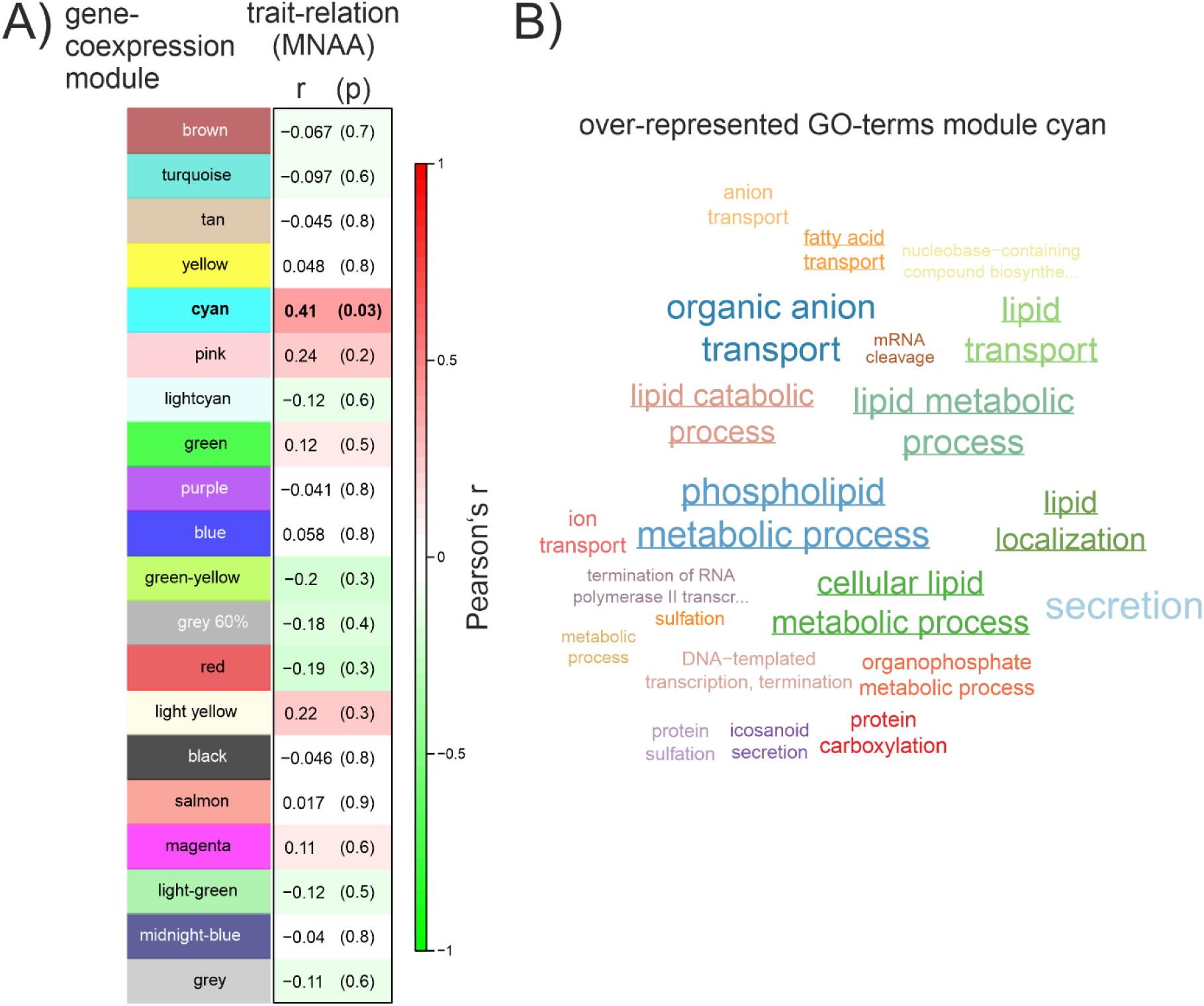
Association between Mean Number of Adaptive Alleles (MNAA) and gene co-expression modules. A) Correlation of MNAA with inferred gene co-expression modules. The modules carry arbitrary colour names. In the right column, the Pearson correlation coefficient between individual transcription levels and the polygenic score and its probability of being identical to zero (in brackets) for the respective module is given. For immediate visual recognition, the correlation coefficient was also translated into a heatmap from green (−1) over white(0) to red (+1). B) Word-cloud of significantly over-represented GO-terms in the cyan gene co-expression module. The font size is proportional to the number of genes. GO-terms with relation to fatty acid metabolism are underlined.

## Discussion

### Natural experiment

In this study, we took advantage of a natural experiment. Following a sampling routine, we coincidently sampled a population pool from a natural *C. riparius* population in late winter just before a cold snap (Pfenninger & Foucault 2020). Such cold snaps at this time of the year are not the rule in Germany, but also not uncommon. The drop of temperature was marked, but not extreme. Likewise, the duration of the snap was relatively short, at least with regard to the temperature drop in the water. It was therefore an event with the potential to leave a selective mark, but it was not an extreme weather event, let alone a catastrophe. This promised the opportunity to study the selective effects of a defined transient event as is typical for the selective regime of fluctuating environments (Bell, 2010). To infer allele frequency changes potentially driven by selection, we therefore sampled another pool from the same site directly after the cold snap.

*C. riparius* does not reproduce at temperatures below 10°C and larval development is nearly stalled. We can therefore rule out selection based on differential reproductive success over the time-span of the experiment (Reznick, 2016). Any potential selection must have occurred in the same generation by differential mortality of different genotypes. At the same time this meant that genetic drift, defined as sampling variance among generations (Wright, 1948), could not have played a role here, since there was no generational passage involved. We have nevertheless included one generation of drift in the calculation of the selection threshold. This threshold is therefore particularly conservative which increases our certainty that the observed allele frequency change beyond this threshold is not due to random processes (Barghi et al., 2019; Hohenlohe, Phillips, & Cresko, 2010). On the other hand, we have certainly missed smaller, yet selection driven allele frequency changes.

### Signs of selection in the natural population

In total, ten regions in the genome showed signs of selection in the data. Given our rather conservative threshold, which required a large change in allele frequency (∼0.5) for detection, we assume that many loci with less pronounced changes remained undetected. The selected haplotypes were very short, often a single SNP, which indicated that they are relatively ancient polymorphisms long since segregating in the population (Nordborg & Tavaré, 2002) and thus separated by recombination from the background in which they arose. Analysis of Tajima’s D for the 1kb windows the selected haplotypes resided in showed that nine out of ten had a positive D, four even in the upper 5% quantile. This indicated that the respective polymorphisms could be regularly under differential, balancing selection (Fijarczyk & Babik, 2015). Given that the presumed selection pressure was a seasonal event, a more or less regularly fluctuating environment with opposite selection pressures in winter and summer appeared plausible. This can lead to a long-term maintenance of the polymorphism under biologically plausible scenarios (Wittmann, Bergland, Feldman, Schmidt, & Petrov, 2017). Recent works suggested that balancing selection could be more widespread than previously thought (Gloss & Whiteman, 2016).

The view that seasonal fitness is related to the different alleles at the identified loci was confirmed by the observation that allele frequencies at the potentially selected loci, with one exception (scaffold 227), returned to near their original frequencies a few generations later. This remarkable correlation between the temporal allele frequency trajectories suggested that the same selection pressure with changing signs among seasons was acting. Seasonally selected polymorphisms with correlated allele frequency trajectories were also observed in natural populations of another dipteran species, *Drosophila melanogaster* (Bergland, Behrman, O’Brien, Schmidt, & Petrov, 2014; Croze et al., 2017). But also the selection-driven beak variability of Galapagos finches in response to different weather conditions in different years, taking into account the different generation times, are a classic example of very rapid evolutionary adaptations to a variable environment (Boag & Grant, 1981). Interestingly, we found a strong negative correlation between the start frequency of the selected SNPs and the absolute deviation from neutrality as measured by Tajima’s D (Supplemental Figure 2). While theory predicts balanced allele frequencies for overdominance (i.e. by heterozygote advantage (Slatkin & Muirhead, 1999)), we are not aware of predictions for expected allele frequencies due to balancing selection by temporally changing selection pressures.

### Validation experiment

We observed a substantial variation in survival time in the validation experiment. Compared to the field observations, it took quite a long time (15 days) until the first larvae started to die. In the field, the temperature dropped for two days only and this was obviously long enough to trigger a substantial mortality. This discrepancy could have several, mutually not exclusive explanations.

First, the lab population had already quite high allele frequencies at the loci in question. If these loci were indeed responsible for the longer survival, the lab population could have been a priori better protected against the cold exposure. This shift in allele frequency relative to the natural population might be due to random drift in the relatively large but nevertheless demographically necessarily restricted lab population. However, the high allele frequencies could also be a tribute to the practice of storing egg ropes at 4°C for a few days prior to initial population set-up and experiments to synchronise their development (Foucault et al. 2019). This could have involuntarily preselected the lab population.

Second, the larvae in the experiment came from normal development at benign temperatures (∼20°C) and were well-fed before they were exposed to 4°C. The larvae in the field likely hatched in autumn and had passed already several months at about 10°C before the cold snap set in. The level of internal resources the two groups could draw upon were therefore likely very different and led thus to mortality faster in the natural population. Lastly, it is likely that the lab experiment did not cover all selection factors that were acting in the field. Given the reduced experimental environment, it is almost certain that the set of selection factors acting on the larvae in the field was different from those in the experiment (Pfenninger & Foucault, 2020a).

The validation experiment confirmed nevertheless the hypothesis that the rising alleles in the natural population did so due to the prolonged cold exposure. This same selection pressure triggered an increase in frequency of the same alleles in the experiment by differential mortality as was observed in nature. This is strong evidence that the candidate alleles indeed played a role in the selection process.

The positive correlation between a straightforward polygenic score (mean number of adaptive alleles per locus involved) and the survival time strongly indicated a relatively simple relation: the more of these alleles were present in an individual, the better were the chances of its longer survival under cold stress. Given the likely involvement of more, but yet unidentified loci, nonlinear interactions among loci and non-quantified environmental components (Sella & Barton, 2019), the degree of determination found here appeared quite substantial. Additional evidence that these loci are co-selected by the same selection pressures came from a strong temporal covariance in allele frequencies, also over longer time scales (Suppl. Figure 3).

### Single locus and multi locus genotype associations with gene transcription data

None of the putatively selected SNPs was within the coding region of an annotated gene. We therefore expected phenotypic effects rather due to changes in the transcription regulation of spatially more or less proximate genes than in structural protein changes.

In an attempt to link the identified loci with basal phenotypic aspects, we sequenced a set of non-lethally cold-exposed individuals for both the genome and the transcriptome. We found transcripts for a substantial proportion of the annotated genes. This is similar to results found in *Drosophila* (Brown et al., 2014). With a what could be called “inversed eQTL approach” (Gilad, Rifkin, & Pritchard, 2008; Majewski & Pastinen, 2011), we explored spatially proximate and thus putative *cis-* interactions between identified selected sites and gene expression levels. By analysing only loci shown to be involved in phenotypic variation and restricting the spatial extent of the search to a plausible range (Schoenfelder & Fraser, 2019), we retained sufficient statistical power even with our relatively limited sample size. Since only two of the identified SNP loci (scaffold227 and scaffold694) showed all possible three genotypes in the sample for which both genotype and transcription data was available, the search for associated cis-regulated genes was necessarily restricted to these.

The gene associated to the selected site on scaffold 227 was identified as Cytochrome P450, family 6, a gene-family that is characteristic for insects (Lewis, Watson, & Lake, 1998). This gene was already several times implicated in the reaction to cold stress(Huang et al., 2017; Lv et al., 2020; Zhang et al., 2015; Zhou, Shan, Tan, Zhang, & Pang, 2019) with an increased transcription rate under cold conditions. It is therefore plausible that an allele associated with an increased transcription rate as observed here is under positive selection under cold stress conditions. While the selected marker SNP on scaffold 227 was spatially closely linked to the gene with the associated transcription regulation, the genes most credibly associated to the SNP genotype on scaffold 694 were both roughly 100 kb up-, respectively downstream. Both genes were also not the closest neighbouring genes, but the 4^th^, respectively 5^th^ transcribed gene up-respectively downstream on the same scaffold. These observations are compatible with recent models of gene-regulation by long range interactions (Schoenfelder & Fraser, 2019) that were also observed in insects (Dorsett, 1999). The gene situated upstream was identified as RFT1 homolog. This protein of the endoplasmatic reticulum membrane appears to be necessary for glycolipid translocation and normal protein N-glycosylation, but its exact function is unknown (Gottier et al., 2017). The associated gene downstream was most similar to a transmembrane receptor protein tyrosine phosphatase DEP1. The selected allele was associated to the downregulation of the respective transcripts. We could not find any studies that have previously linked this gene to cold stress.

Using the polygenic score as a proxy for survival time on the individual level was associated to 28 differentially expressed genes. One third of these (nine) belong to (larval) cuticule proteins, endocuticule proteins or endochitinase, suggesting a role for these genes in differential survival. Moreover, the polygenic score as a proxy for survival time on the individual level revealed a moderately strong correlation (r = 0.4) to a co-expression module. The co-expression module was statistically enriched for genes involved in fatty acid metabolism. The fatty acid metabolism is known to be crucial for the overwintering of insects (Sinclair & Marshall, 2018; Storey & Storey, 2013; Toprak, Hegedus, Doğan, & Güney, 2020), i.e. under cold stress e.g. by providing the necessary energy storage resources or keeping membranes subtle by changing their fatty acid composition (Overgaard, Sørensen, Petersen, Loeschcke, & Holmstrup, 2005). There are many examples showing that unsaturated fatty acids increase under cold temperatures in insects (reviewed in Clark and Worland 2008; Teets and Denlinger 2013). Moreover, fat content was one amongst other phenotypes linked to exposure to a new temperature in *Drosophila*, which also appears to be a polygenic trait (Barghi et al., 2019).

The association suggested that at least some of the identified loci are directly or indirectly involved in *trans*-regulation of the transcription of these genes. One of the spatially closest genes to a selected site was a histone acetyltransferase p300 on scaffold673. These genes are high level switches that regulate broad scale gene transcription pattern via chromatin remodelling (Tropberger et al., 2013). Amongst many other biological functions, it has been shown that the gene is responsible for the differentiation of fat cells (Gesta, Tseng, & Kahn, 2007). The transcription level of this gene was significantly associated with the observed genotypes (Supplemental Figure 3), suggesting that a *trans*-regulated altered transcription of this gene could provide a link between the multilocus genotype and the fatty acid metabolism genes.

This finding allowed to hypothesise a particular physiological trait(s) the identified selected SNPs could, amongst others, contribute to the observed variation in survival time. Individuals with a high polygenic score for the loci involved could accumulate more fat reserves, which increased survival time under prolonged cold conditions. This clearly beneficial fitness effect during the winter season could be reversed in summer, when the accumulation of then unnecessary fat reserves deviates resources from reproduction or increases attractivity for predators. Such a more or less regularly fluctuating selection regime on a polygenic trait could well conform to the theoretical preconditions necessary to maintain the respective polymorphisms over longer periods (Wittmann et al., 2017). Whether there is really a link between the extent of the individual fat reserves and the observed survival time under cold stress conditions, however, remains to be tested, just like a causal relation between the respective multilocus genotype and the regulation of the fatty acid metabolism. However, many more intermediate level traits, e.g. changes in cuticule composition likely contributed to survival under constant cold exposure.

## Conclusions

In this study we could show that normal, short term environmental variability can lead to measurable natural selection on a polygenic trait in a natural population. Time series population genomic analyses from field samples obviously have the power to pick up such transient signals, even if they consist rather of moderate to strong changes in allele frequencies than in fixation of loci. The observed high temporal correlation of the allele-frequency changes of the loci involved holds the promise that changes in the selective regime may be identified from population genomic time series data.

## Supplemental Material

**Supplemental Figure 1.**
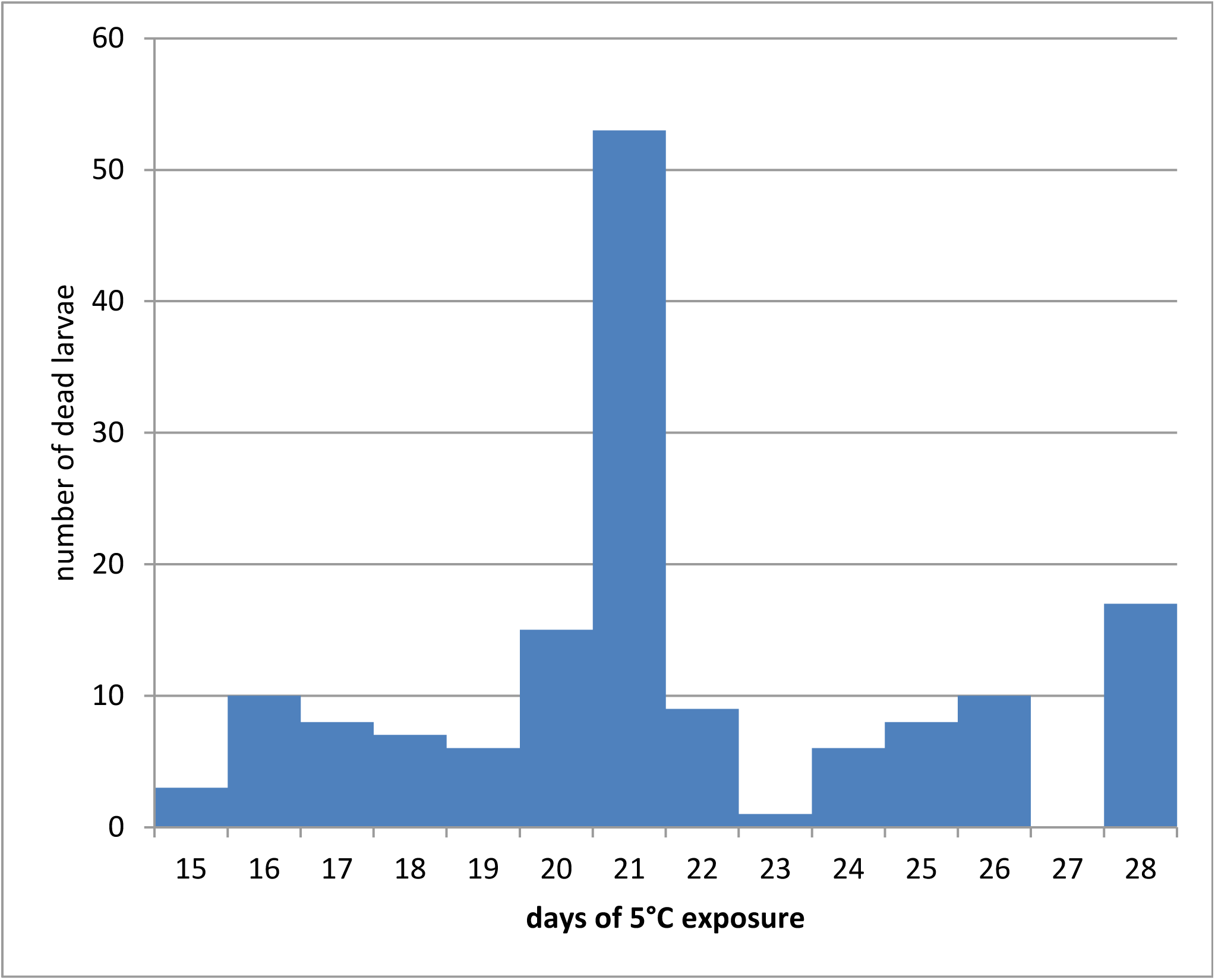
Distribution of dying days in the experimental confirmation. The experiment ended on the 28^th^ day. On this day, 11% of the larvae were still alive.

**Supplemental Figure 2.**
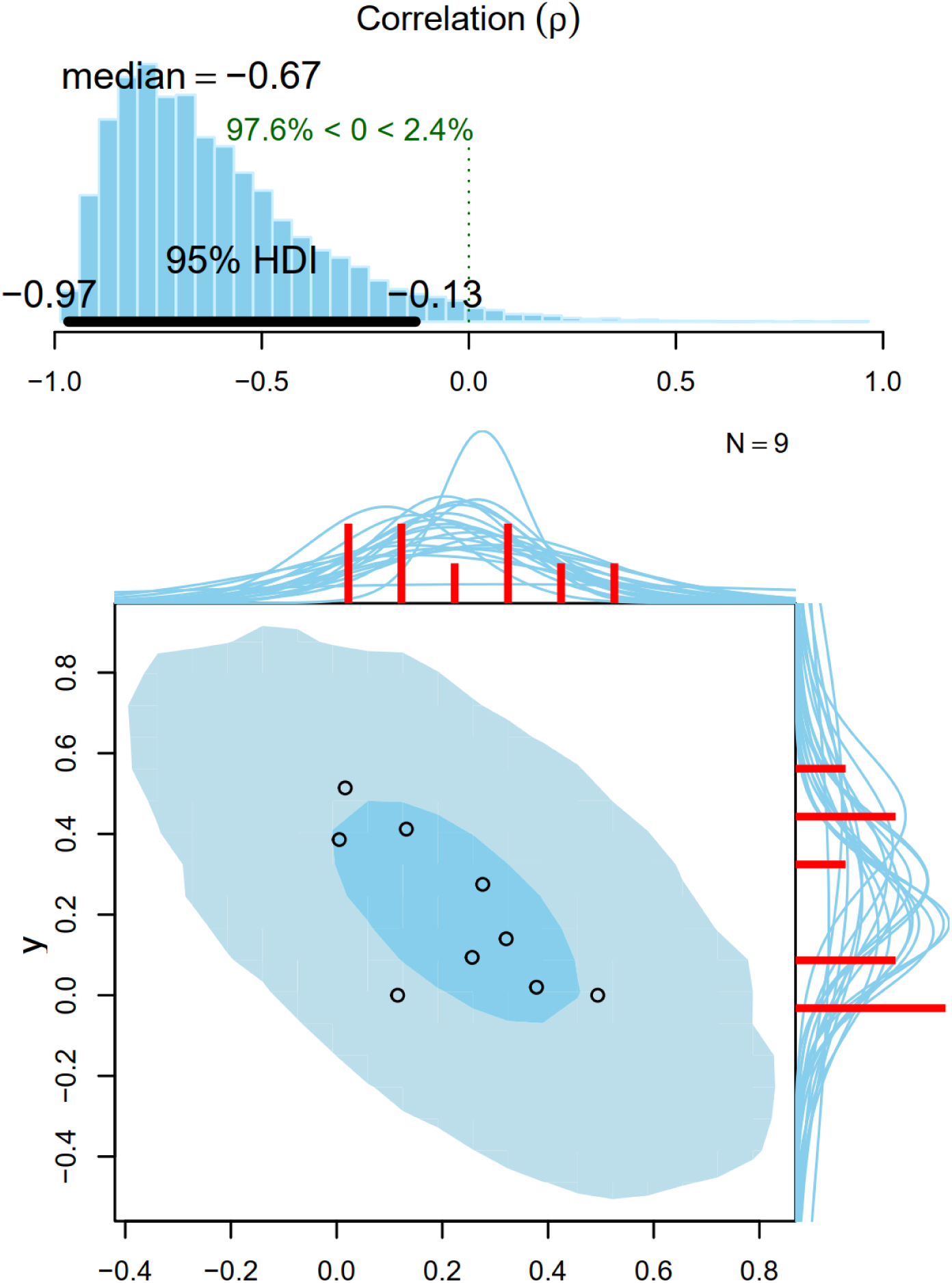
Bayesian analysis of correlation between the starting allele frequency and the absolute deviation from neutrality in Tajima’s D for the 1 kb windows of the selected SNPs.

**Supplemental Figure 3.**
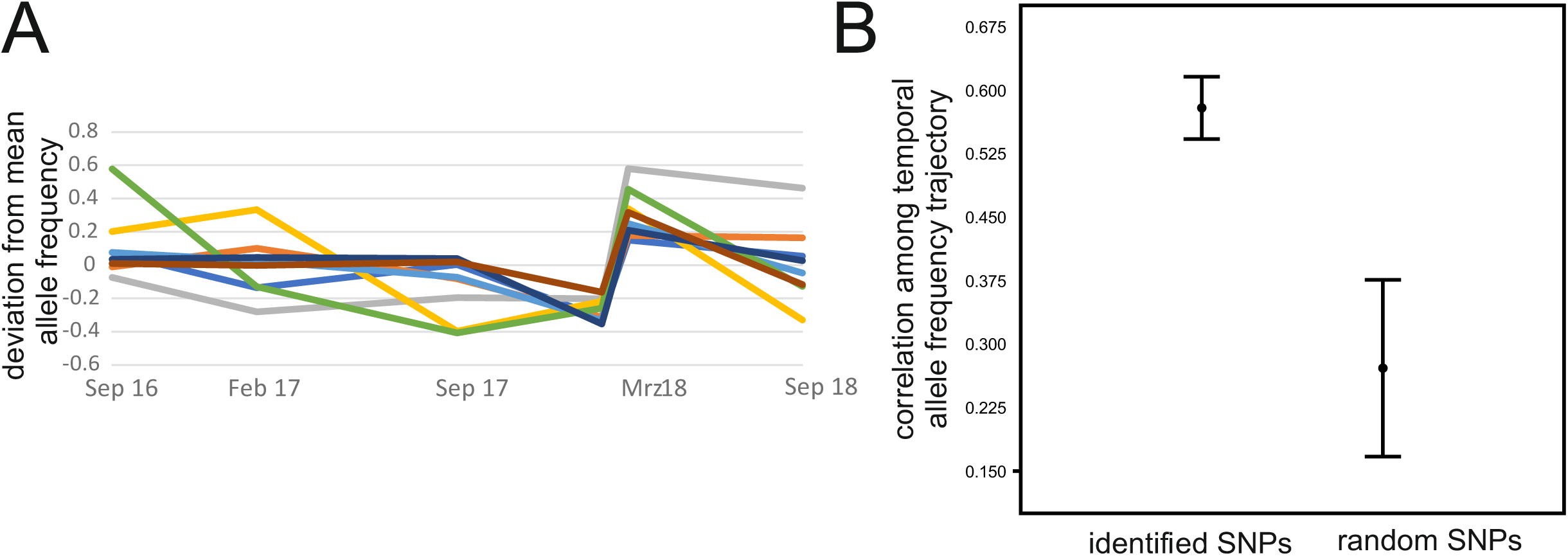
A) Temporal covariation of allele frequencies at identified loci over time. Shown is the deviation from the mean frequency per locus. B) Difference in mean pairwise correlation coefficient compared to a random sample of 10 SNPs with similar variance located on different scaffolds. Data from Pfenninger, M., & Foucault, Q. (2020). Quantifying the selection regime in a natural *Chironomus riparius* population. bioRxiv 2020.06.16.154054.

**Supplemental Figure 4.**
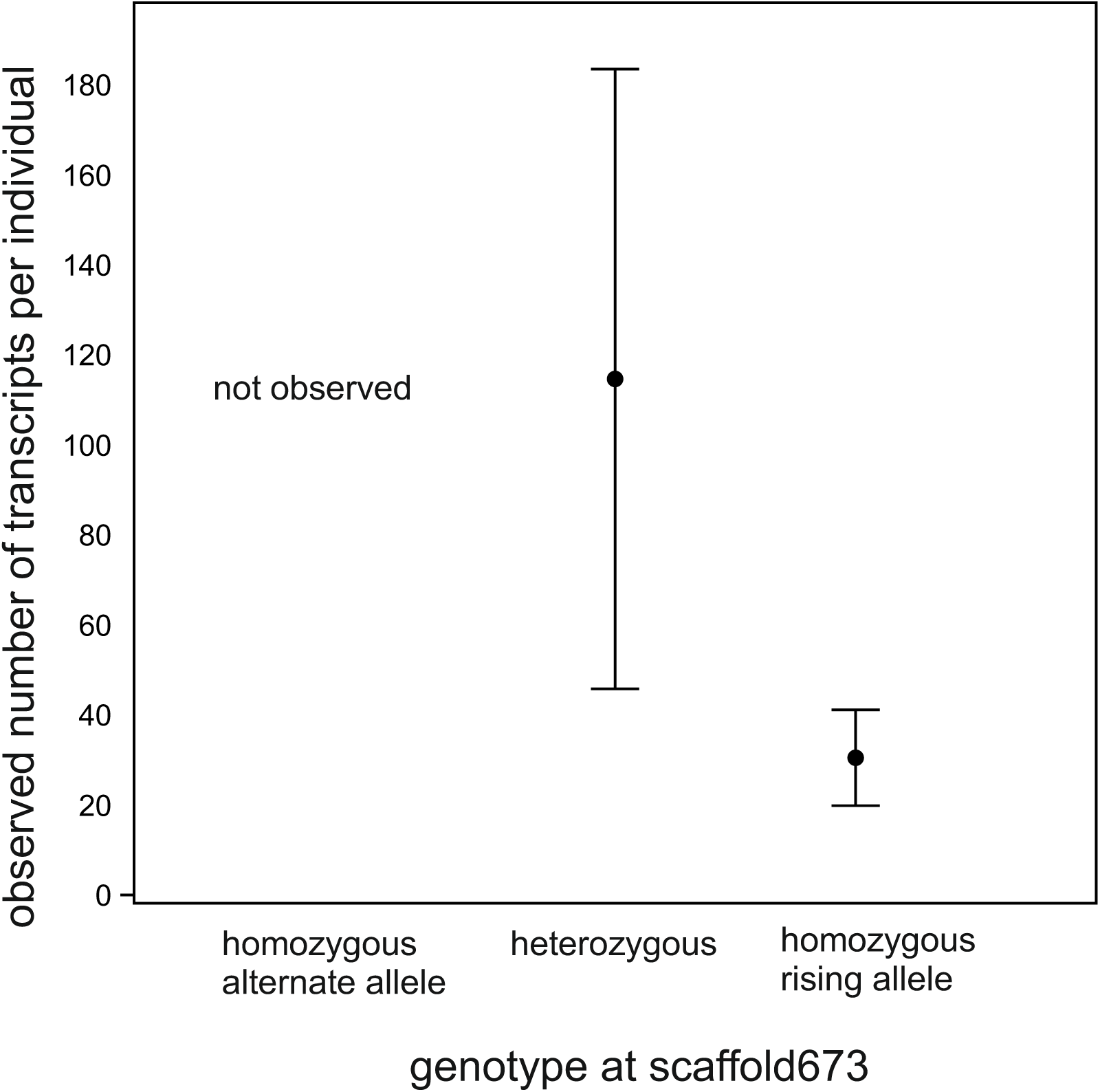
Associations of genotype at scaffold673 site with gene expression levels of the closest gene histone acetyltransferase p300. The difference between the transcription levels among the two observed genotypes was significant (Mann-Whitney test U = 27, p = 0.008).

